# A vesicle microrheometer for high-throughput viscosity measurements of lipid and polymer membranes

**DOI:** 10.1101/2021.03.04.433848

**Authors:** Hammad A. Faizi, Rumiana Dimova, Petia M. Vlahovska

## Abstract

Viscosity is a key property of cell membranes that controls mobility of embedded proteins and membrane remodeling. Measuring it is challenging because existing approaches involve complex experimental designs and/or models, and the applicability of some is limited to specific systems and membrane compositions. As a result there is scarcity of data and the reported values for membrane viscosity vary by orders of magnitude for the same system. Here, we show how viscosity of bilayer membranes can be obtained from the transient deformation of giant unilamellar vesicles. The approach enables a non-invasive, probe-independent and high-throughput measurement of the viscosity of bilayers made of lipids or polymers with a wide range of compositions and phase state. Pure lipid and single-phase mixed bilayers are found to behave as Newtonian fluids with strain-rate independent viscosity, while phase-separated and diblock-copolymers systems exhibit shear-thinning in the explored range of strain rates 1-2000 *s*^−1^. The results also reveal that electrically polarized bilayers can be significantly more viscous than charge-neutral bilayers. These findings suggest that biomembrane viscosity is a dynamic property that can be actively modulated not only by composition but also by membrane polarization, e.g., as in action potentials.

---

Cells and cellular organelles are enveloped by membranes, whose main structural component is a lipid bilayer [1]. The lipid bilayer endows membranes with fluidity that is essential for functions that depend on biomolecules mobility, e.g., signaling [2–4]. Fluidity is modulated by membrane composition and this homeoviscous adaptation is crucial for the survival of organisms that can not regulate their body temperature like bacteria [5, 6]. Viscosity is the common measure for fluidity, yet for membranes this property has been challenging to assess. Data for viscosity of lipid membranes are limited and reported values vary significantly, sometimes by orders of magnitude for the same system (SI Table 1). For example, reported values for the surface shear viscosity of membranes made of a typical lipid such as dioleoylphosphatidylcholine (DOPC) span two orders of magnitude: (0.197 *±* 0.0069) *×* 10^−9^ Pa.s.m [7], (1.9 *±*11) *×* 10^−9^ Pa.s.m [8], (16.72 *±* 1.09) *×*10^−9^ Pa.s.m [9]. For a similarly structured lipid, palmitoylphosphatidylcholine (POPC), the surface viscosity measured by shear rheology of Langmuir monolayers is 3 *×* 10^−4^ Pa.s.m [10]. Experimental methods that utilize free-standing bilayer membranes, e.g, vesicles or black lipid membranes, rely on estimates from the rate of tether formation [11], diffusion coefficients of domains [12–14] or membrane-anchored nanoparticles [15, 16], domain shape fluctuations [17], domains motion on vesicles induced by applied flow [8, 18], bilayer thickness fluctuations or lipid dynamics measured with neutron spin echo spectroscopy [19– 21], fluorescence quantum yield or lifetime of viscosity-sensitive fluorescent dyes [22, 23], and the forced motion of colloidal particles in the membrane [24–26]. In silico approaches, using molecular dynamics simulations, have also been developed to determine membrane viscosity [7, 27, 28]. Despite these advances, the systematic study of membrane viscosity has been hindered by various limitations of the proposed methodologies. For example, domain-based methods [8, 12, 13, 17, 18] are limited to phase-separated membranes and the measured viscosity reflects the continuous phase, not the membrane as a two-phase fluid. Bilayer thickness fluctuations [19] depend on both shear and dilational monolayer viscosities. In particle-based methods [15, 16, 24–26], the probe perturbs the membrane and the data interpretation requires complicated analysis that discerns the contributions to the particle mobility from the flow in the membrane and the surrounding fluids [29–31]. Furthermore, since membrane surface viscosity is a macroscopic quantity, defined on scales where the bilayer can be modeled as a two-dimensional incompressible fluid, methods utilizing measurements at the micro- or nano-scale and/or based on molecular probes may not report the effective continuum viscosity but a quantity, often called “microviscosity”, which is local and depends on the immediate environment [21]. These complexities are likely the source of the huge variability in reported values of viscosity for lipid bilayer membranes, making it challenging to compare data obtained by different methods.

**TABLE I.** Membrane viscosity and bending rigidity for various bilayer systems at 25.0 ^*o*^C and *E*_0_ = 8kV/m (strain rate 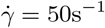).

| Composition | Viscosity<br>(nPa.s.m) | Bending Rigidity<br>($k_B T$) |
| --- | --- | --- |
| DOPC | $4.11 \pm 2.63$ | $22.2 \pm 2.0$ |
| OMPC | $7.73 \pm 3.09$ | $27.1 \pm 2.6$ |
| POPC | $9.32 \pm 5.95$ | $27.8 \pm 2.3$ |
| SOPC | $9.73 \pm 5.80$ | $30.1 \pm 3.1$ |
| DOPC:Chol | $7.00 \pm 4.77$ | $27.8 \pm 4.6$ |
| DPPC:DOPC:Chol (1:1:1) | $17.7 \pm 3.06$ | $72.0 \pm 8.4$ |
| DPPC:DOPC:Chol (1:1:2) | $15.4 \pm 2.40$ | $69.2 \pm 7.9$ |
| DPPC:Chol (1:1) | $56.4 \pm 4.63$ | $121.3 \pm 11.0$ |
| PBd<sub>13</sub>-*b*-PEO<sub>11</sub> | $14.4 \pm 4.40$ | $17.1 \pm 1.5$ |
| PBd<sub>22</sub>-*b*-PEO<sub>14</sub> | $686 \pm 51.0$ | $31.0 \pm 5.1$ |
| PBd<sub>33</sub>-*b*-PEO<sub>20</sub> | $2890 \pm 670$ | $54.4 \pm 6.4$ |
| PBd<sub>46</sub>-*b*-PEO<sub>24</sub> | $20600 \pm 4700$ | NA |
| PBd<sub>54</sub>-*b*-PEO<sub>29</sub> | $46700 \pm 900$ | $154.0 \pm 16.0$ |
| PBd<sub>120</sub>-*b*-PEO<sub>78</sub> | $157000 \pm 54500$ | NA |

Here, we show that the deformation of giant unilamellar vesicles (GUVs) can be employed to obtain the surface “macroviscosity”, i.e., the shear viscosity of the membrane treated as a two-dimensional incompressible fluid. Upon application of an extensional stress, e.g., generated by an uniform electric field [32–35], extensional flow [36–38], or an optical stretcher [39, 40] a quasi-spherical vesicle deforms into a prolate ellipsoid. The aspect ratio, *ν* = *a/b* (see sketch in Fig. 1A), increases and reaches a steady state. When the stress is removed, the vesicle relaxes back to its equilibrium spherical shape, see Fig. 1B. The reproducibility of the results was tested by repeated measurements with the same vesicle, see Fig. 1B inset, showing identical slope of the aspect ratio curves for small deformations.

**FIG. 1.**
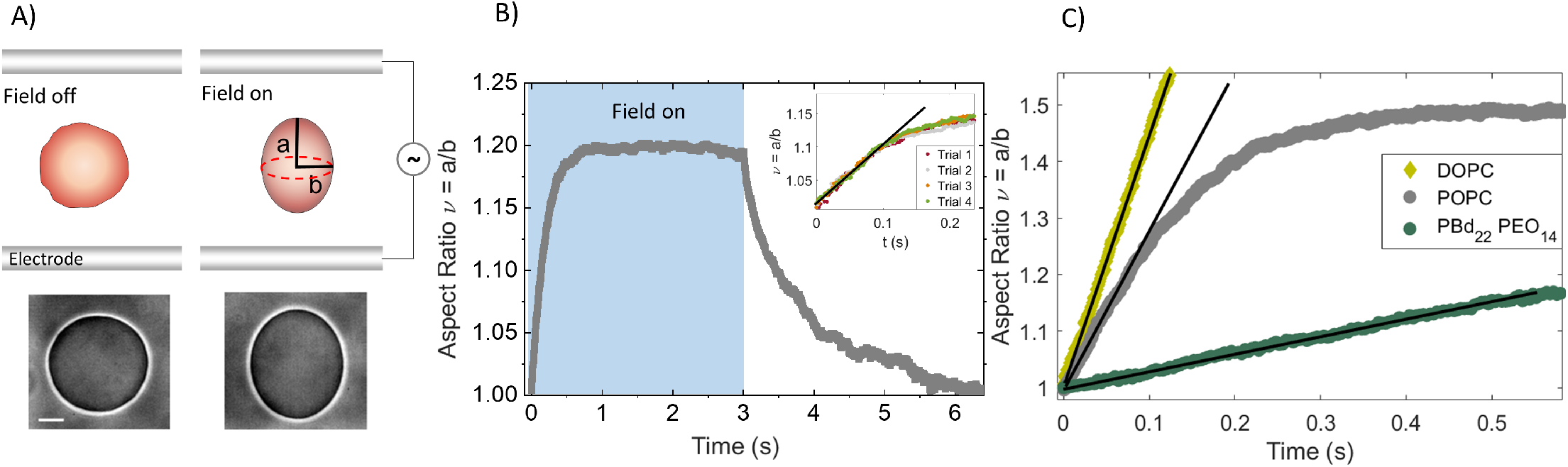
Electrodeformation method to measure membrane viscosity. (A) A uniform electric field deforms a GUV into a prolate ellipsoid by pulling out area stored in suboptical thermally-excited membrane undulations. Snapshots of the vesicle during the experiment. Imaging with phase contrast microscopy. Scale bar: 15 *µ*m. (B) Prolate deformation of a POPC GUV in an electric field with amplitude *E*_0_= 10 kV/m and frequency of 1 kHz. Time zero in all graphs corresponds to turning the field on.The inset shows that repeated deformation does not alter the initial slope of the deformation curve. C) Vesicles made of lipids (DOPC, POPC) and the diblock-copolymer PBD_22_PEO_14_ deform at a different rate indicating different membrane viscosity. The field strength and frequency are 8 kV/m and 1 kHz. The solid lines correspond to the theoretical fit with Eq. 2.

Even though the applied stress is extensional and vesicle deformation is axisymmetric, material transported on the vesicle surface undergoes shear because the membrane is area–incompressible [41–45]. The rate at which the vesicle elongates while the field is on, and relaxes back to its equilibrium shape after the field is turned off, is related to the membrane shear viscosity. For small deformations, *ν* ≲ 1.3, the evolution of the aspect ratio is described by [34, 45–47] (see Appendix Section 2 for a summary of the theory)

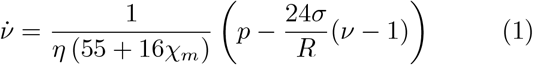

where *χ*_*m*_ = *η*_*m*_*/ηR* is the dimensionless surface viscosity *η*_*m*_, *η* is the viscosity the solution inside and outside the vesicle (assumed to be the same), *σ* is the membrane tension, and *R* is the vesicle radius. In an applied extensional flow with strain rate 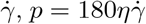. In the case of a charge-neutral vesicle in a uniform DC electric field with amplitude *E*_0_, 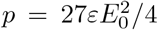, and in an AC field with frequency *ω, p*(*ω*) is given in Appendix Section 2. Thus from the vesicle dynamics in response to an extensional flow or a uniform electric field it is straight-forward to obtain the membrane viscosity. Note that the dynamics does not involve dilational viscosity because vesicle deformation and the accompanying increase in apparent area comes from ironing of suboptical thermally-excited membrane undulations, while the area per lipid remains the same.

## RESULTS

We illustrate the implementation of the approach on the example of a quasi-spherical vesicle subjected to a uniform AC electric field. The electric field is advantageous over extensional flow or radiation pressure in optical stretchers because of (i) the simplicity of the experimental set up, and (ii) the ability to create potential difference across the bilayer emulating transmembrane potentials in living cells. The use of an AC field is preferable because (i) a DC field could cause Joule heating and electroosmotic flows, whose effect on vesicle deformation and stability is difficult to account for [48], and (ii) the transmembrane potential can be modulated by changing the electric field frequency [49, 50]. We applied the method to fluid membranes composed of the phosphatidylcholine lipids palmitoyloleoyl (POPC), dioleoyl (DOPC), oleoylmyristoyl (OMPC), stearoyloleoyl (SOPC), and dipalmitoyl (DPPC), cholesterol (chol) and diblock copolymers, PBd_*x*_-*b*-PEO_*y*_ with varying hydrophobic molecular weight, M_*h*_, from 0.7 kDa to 6.8 kDa.

### Vesicle transient deformation yields membrane viscosity

Figure 1 summarizes the experiment. The elongation curves of a GUV initially show linear increase (Figure 1C and Appendix Figure S3). The linear slope is predicted by Eq. B8 if the second term, which describes the action of the tension opposing the deformation, is neglected

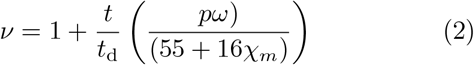

where 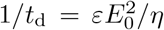 is the characteristic rate-of-strain imposed by the electric field. Thus, the slope of the aspect ratio plotted as a function of the rescaled time *t/t*_d_, at the same field frequency, depends solely on the membrane viscosity. Vesicles made of different lipids or diblock-copolymers show different initial slopes, see Fig. 1C, indicating different membrane viscosities. The linear response is observable only if the restoring force of the membrane tension is negligible compared to the deforming electric stress, i.e., 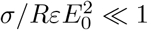, which is indeed the case for typical values of the applied electric field *E*_0_ = 5 kV/m, equilibrium membrane tension *σ* = 10^−8^ N/m, and vesicle radius *R* = 10 *µ*m. The time up to which the linear approximation is reasonable (see Appendix Section 2B) is estimated to be 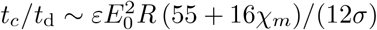. Using times up to 0.1*t*_*c*_*/t*_d_ (or equivalently *ν* up to 1.1) minimizes the error in the linear fit.

### Viscosity shows dependence on field amplitude and frequency

The stress, generated by the electric field, shears the membrane with a characteristic rate 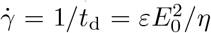. Modulating the field amplitude thus enables us to vary the shear rate in a wide range and to probe if bilayers behave as Newtonian fluids: with shear-rate independent viscosity. Increasing *E*_0_ from 1 to 50 kV/m at a given frequency increases the effective shear rate from 1 *s*^−1^ to 2000 *s*^−1^. We find that bilayers made of only one lipid or a homogeneous mixture, either in the liquid disordered (e.g., DOPC or POPC) or liquid ordered (e.g., DPPC:Chol 1:1) state, exhibit rate-independent viscosity (Fig. 2A). Phase-separated bilayers such as DPPC:DOPC (1:1), which have solid domains coexisting with liquid-disordered continuous phase (see inset of 2B) [51–53], and the diblock-copolymer membranes shear thin, i.e., their viscosity decreases with increasing shear rate (Fig. 2B-C and Appendix Section 3).

**FIG. 2.**
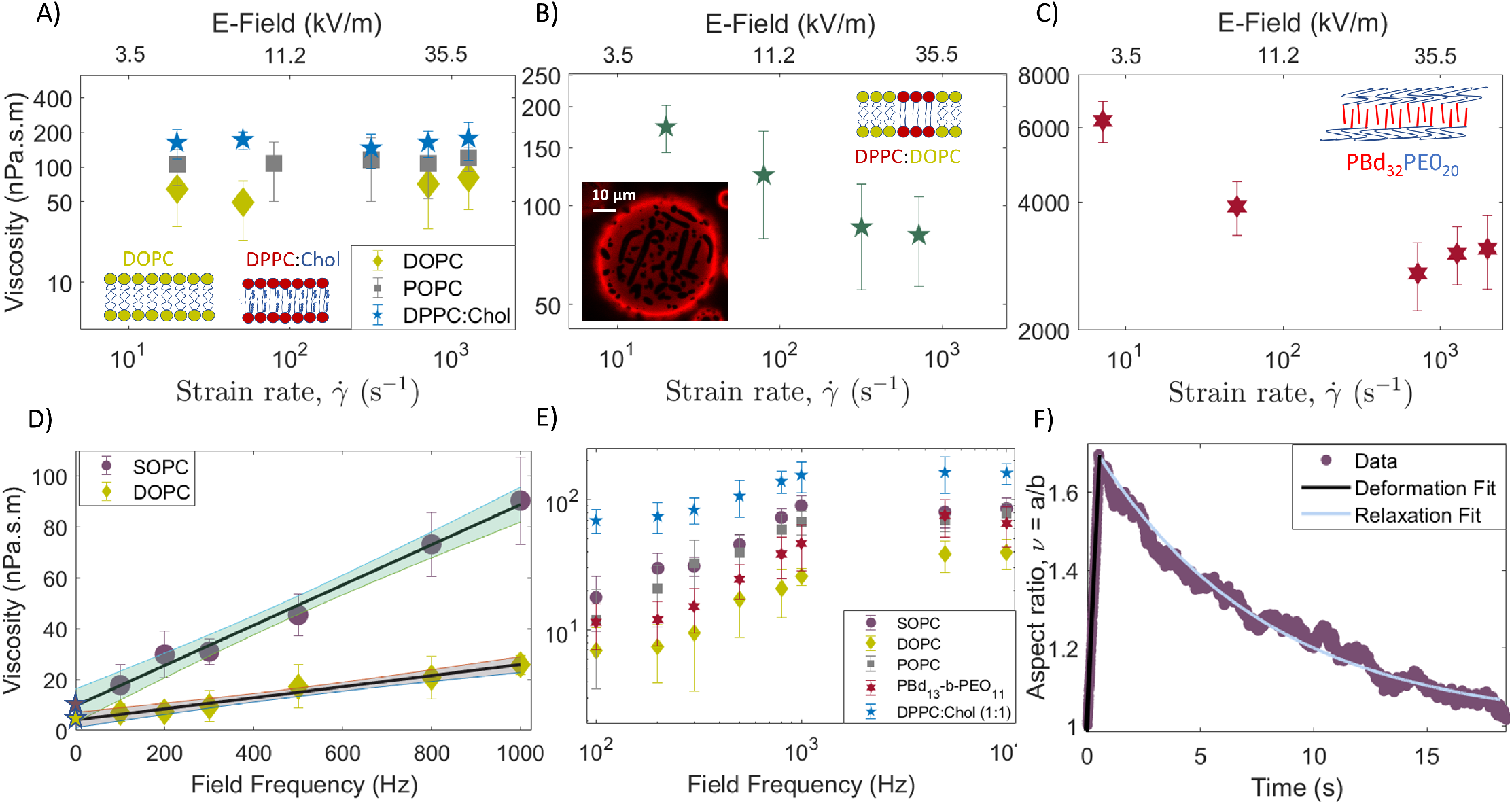
Viscosity dependence on electric field strength, or equivalently strain rate 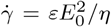, and frequency. A) Single-component (DOPC or POPC) and single-phase multicomponent bilayers (DPPC:Chol 1:1) behave as Newtonian fluids. B) Viscosity of the phase-separated multicomponent (DPPC:DOPC 1:1) decreases with the effective strain rate. The inset shows an intricate network of finger-like domains indicative of fluid (red) and gel (dark) phase coexistence. C) Diblock-copolymer bilayers also display shear thinning viscosity. All measurements in panels A-C, were done at field frequency 1kHz. D) Viscosity measured at different frequencies at a fixed strain rate 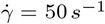, corresponding to field strength 8 kV/m, increases with frequency. Extrapolating to zero frequency (the DC limit) yields the shear surface viscosity. The symbols on the vertical-axis-intercept refer to the extrapolated values of 4.11*±* 2.63 nPa.s.m and 9.73 *±*5.80 nPa.s.m for DOPC and SOPC, respectively. The R-squared value of the linear fit is 0.98 and 0.97 for the SOPC and DOPC, respectively. The shaded band represents a 95% confidence interval. E) The viscosity reaches as plateau at frequencies above few kHz. This behavior is common to all studied compositions. F) Shape relaxation after the field is turned off yields the similar value for the viscosity as the zero-frequency limit obtained from the frequency sweep. Data for SOPC, *E*_0_ = 8 kV/m and *ω* = 1 kHz. Tension obtained from flickering spectroscopy is 5.7 *±* 2.6 *×*10^−9^ N/m. Viscosity obtained from initial deformation and relaxation is 164 nPa.s.m and 21.1*±* 29.4 nPa.s.m, respectively.

Intriguingly, measurements at the same field amplitude but different frequencies revealed an apparent increase of viscosity with frequency (Fig. 2D). More precisely, the viscosity shows an initial linear increase with frequency and a plateau above few kHz (Fig. 2E). The behavior is observed with all studied compositions. Viscosity values in the literature obtained by methods that do not involve electric fields (Appendix Table II) are at least a factor of 10 smaller than the plateau-value of the viscosity but comparable to the zero-frequency limit obtained from the linear extrapolation at low frequencies. We hypothesize that the zero-frequency viscosity is representative for the viscosity of the membrane in the absence of electric field. As a test, we analyzed the vesicle relaxation back to its equilibrium shape, after the field is turned off (Fig. 2F). That analysis, however, is complicated by the fact that unlike the initial elongation upon application of the field, the relaxation is driven by and thus depends on the membrane tension. In order to leave the membrane viscosity as the only fitting parameter for the relaxation curve, the tension needs to be independently determined. In this experiment, the tension was obtained from the analysis of the equilibrium thermally-excited membrane undulations (flickering spectroscopy) in the absence of electric field before the electrodeformation experiment. The fit of the relaxation curve with Eq. B8 yielded viscosity 21.1 *±* 29.4 nPa.s.m. The large error is due to the uncertainty in the tension. The viscosity value is close to the zero-frequency limit of the viscosity obtained from the frequency sweep (9.73 *±* 5.80 nPa.s.m) for SOPC. We adopt the frequency sweep to determine the initial linear dependence of the viscosity and its “no-field” limit.

To summarize, the method involves measuring apparent viscosities at different frequencies in the range 0.1-1 kHz and extrapolating to zero-frequency (as in Fig.2D) to obtain the value of the viscosity in the absence of electric field. Electric field of 8 kV/m (strain rate 50 *s*^−1^) produces a good range of data in the initial linear deformation regime. Measurement and data analysis of one vesicle typically takes about 10 min. We analyze 10-50 vesicles per viscosity value. The viscosity obtained for 14 different bilayer compositions is reported in Table I. The values are in the range reported in previous studies (Appendix Table II). For example, the bilayer viscosity for SOPC is in good agreement with previously reported values from 3 to 13 nPa.s.m [26, 27, 54].

### Viscosity correlation with lipid diffusivity, membrane composition and thickness

Since mobility of lipids or domains is often used to asses membrane fluidity [12–14], we have compared our results for membrane viscosity with measurements of the diffusion coefficient of a lipid dye (DiI-C18) using Fluorescence Correlation Spectroscopy (FCS) in the same ternary system [55]. The diffusion coefficient scales inversely with surface viscosity (Fig. 3), a trend expected from the Saffman-Delbrück’s model [56], *D* = *k*_*B*_*T/*(4*πη*_*m*_) (log(*η*_*m*_*/*(*ηr*)) − 0.5772). However, there is a quantitative disagreement: using lipid dye radius *r* = 0.5 nm (comparable to the radius of DOPC estimated from the area per lipid head, 67.4-75.4 °*A*^2^ [57–62]) the Saffman-Delbrück’s equation predicts much lower diffusivities. These results suggest that while increasing viscosity does correlate with decreasing diffusivity, it is not trivial to relate membrane viscosity and the diffusion constant because diffusion of molecular probes is sensitive to the probe itself as well as the bilayer structure [11, 14, 21]. However, the empirically obtained dependence in Fig. 3, could presumably be used to roughly deduce membrane viscosity from molecular diffusivity measurements.

**FIG. 3.**
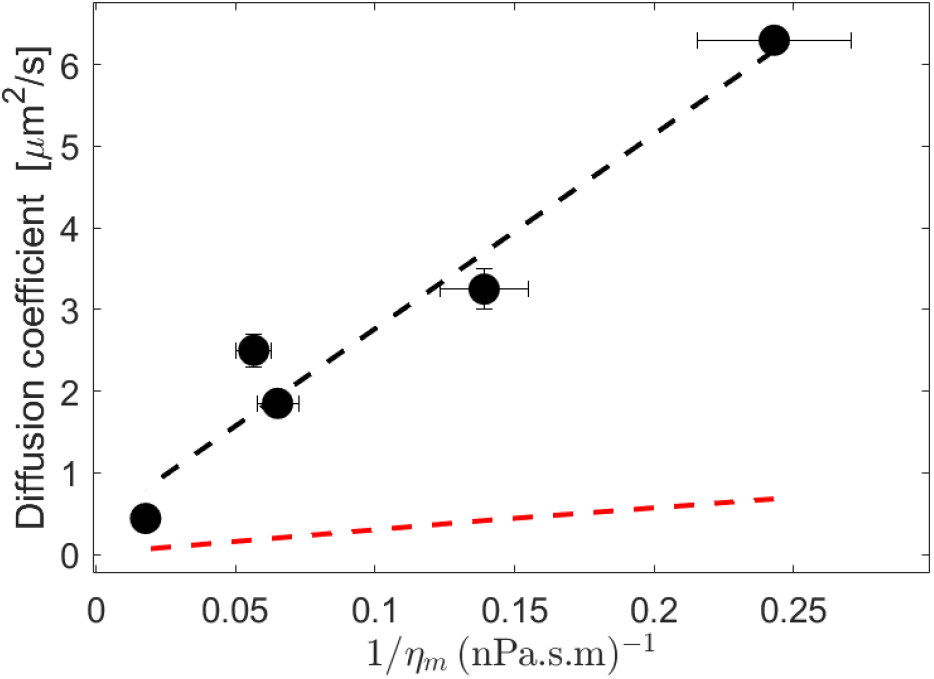
Membrane viscosity as a function of diffusivity values obtained with FCS *Scherfeld et al*. [55] for membrane compositions DOPC, DOPC:Chol (1:1), DPPC:DOPC:Chol (1:1:2), DPPC:DOPC:Chol (1:1:1) and DPPC:Chol (1:1). Values of the diffusion coefficient are listed in the SI. The red dashed line corresponds to the prediction from the Saffman-Delbrück’s model with probe radius *r* = 0.5 nm. The black dashed line is a linear fit with intercept 0.383 *µ*^2^m/s and slope 23.83 *µ*m^3^.mPa

The viscosity values complied in Table I highlight that in pure lipid systems membrane viscosity decreases with the number of unsaturated bonds in the hydrophobic tail. POPC, OMPC and SOPC have a single unsaturated bond, while DOPC has two double bonds in the hydrophobic tails. DOPC bilayers exhibit much smaller viscosity than the other single unsaturated lipid bilayers. In the mixed systems, adding DPPC or/and cholesterol to DOPC increases the viscosity. Adding cholesterol (molar ratio 1:1) to DOPC increases the bilayer viscosity as cholesterol increases the packing of the liquid disordered (*L*_*d*_) phase. For liquid ordered (*L*_*o*_) phase, such as DPPC:Chol (1:1), the bilayer viscosity is much higher due to the tight packing provided by saturated acyl chains. The effect is less pronounced in the ternary *L*_*o*_ system DOPC:DPPC:Chol (1:1:2) and DOPC:DPPC:Chol (1:1:2). For a phase-separated system of coexisting solid and fluid phases DOPC:DPPC (1:1), see Fig. 2B, the viscosity also increases relative to pure DOPC. The increase agrees with an estimate of the viscosity of a 2D suspension, *η*_*eff*_ = *η*_*DOP C*_(1 + 2*ϕ*), where *ϕ* ∼ 0.4 is the fraction of the solid phase [63].

Using our method we were also able to examine the commonly assumed relation between the 2D (*η*_*m*_) and 3D viscosities (*η*_3*D*_), *η*_*m*_ = *η*_3*D*_*h*, where *h* is the membrane thickness. The thickness of diblock-polymer bilayers increases with the molecular weight of the hydrophobic part, 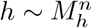 [64, 65], *where n* lies within the theoretical bounds of 0.5 (random Gaussian coil) and 1 (full stretch). The polymers’ molecular weight varies in a wide range (1-8 kDa) thereby resulting in bilayers with greater range of thicknesses, unlike lipids. The diblock polymers showed membrane viscosity spanning four orders of magnitude range, from 14 nPa.s.m to 157000 nPa.s.m. The lowest molecular-weight polymer membrane exhibits a viscosity similar to that of POPC. The membrane viscosity in Fig. 4A does follow a power-law dependence on *M*_*h*_, but the power 5.6 is much larger than the expected range 0.5-1. In contrast, as seen from Fig. 4B, the bending rigidity follows a power-law consistent with the expected 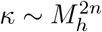, since *κ/K*_*A*_ ∼ *h*^2^, where *K*_*A*_ is the area-compressibility modulus [66]. The *K*_*A*_ value is relatively insensitive to the molecular weight; its value for the lipids was taken to be 250 mN/m and for the polymer membranes - 100 mN/m [67, 68]. The polymer 3D viscosity *η*, however, varies with the polymer molecular weight and this may be the source of the unexpectedly higher exponent in the power-law dependence of *η*_m_ on *M*_*h*_. Comparison of the dependence on the strain-rate of the 2D and 3D viscosities (Fig. 4C and Appendix Section S4) shows that in three dimensions the shear-thinning is more pronounced. This suggests that the polymer confinement into a bilayer impedes the microstructure dynamics responsible for the shear thinning. Intriguingly, taking the ratio of the 2D and 3D viscosities yields a thickness *h* ranging from 200 to 350 nm much larger than typical thickness measured with cryo-TEM [69]. The disparity might arise from the thermally-driven membrane undulations, which result in interface roughness that is in the order of 100 nm for a tension-free membrane and 20 *k*_*B*_*T* bending modulus. Thus the effective membrane thickness would be larger for ‘fuzzy’ membranes.

**FIG. 4.**
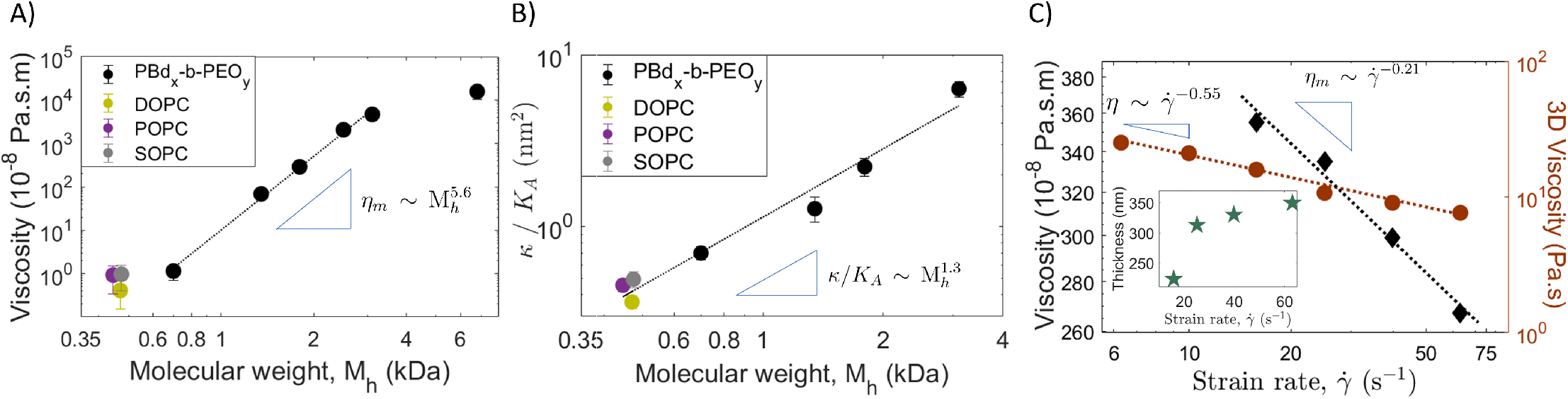
A) Viscosity of PBd_*x*_-*b*-PEO_*y*_ bilayers as a function of the *M*_*h*_ of the hydrophobic part (PBd). Viscosities of phospholipids are also shown for comparison. B) Bending rigidity of PBd_*x*_-*b*-PEO_*y*_ and phospholipids bilayer membranes as a function of molecular weight. The power law obtained is 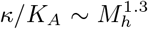. C) Comparison of the shear-thinning of the 2D and 3D viscosities of PBd_33_-*b*-PEO_20_. The inset shows the membrane thickness obtained from *η*_*m*_*/η* = *h*.

## DISCUSSION AND CONCLUSIONS

We developed a method to measure the shear viscosity of the two-dimensional bilayer fluid using the transient deformation of a giant vesicle. The approach is inspired by the interfacial rheology measurements using deformation of droplets and capsules [70]. The “vesicle-rheometer” enabled the collection of an unprecedented amount of data, which provided valuable insights into the rheology of bilayer membranes and lead to two new intriguing findings that raise more questions.

First, bilayer membranes can behave as non-Newtonian fluids. Phase-separated bilayers are the 2D analogs of 3D dispersed systems such as emulsions and suspensions, which exhibit shear thinning behavior due to microstructure reorganization and relaxation; the domain dynamics in the phase-separated bilayers is likely the source of the shear thinning dynamics. Likewise, the shear thinning viscosity of the diblock-copolymer membranes likely originates from polymer stretching and relaxation on a time scale comparable to the shearing.

Second, membrane viscosity increases with field frequency reaching a plateau above few kHz. The zero-frequency limit of the viscosity is comparable to the one measured in the absence of electric field by other methods, thus the frequency effect is not related to the electric field amplitude. We hypothesize that the phenomenon is due to membrane electrical polarization. In an applied electric field, ions brought by conduction accumulate at the membrane surfaces, because the bilayer is impermeable to ions (Fig. 5). At low frequencies, *ω* ≪ *ω*_*c*_ *∼ λ*_v_*/RC*_*m*_, where *C*_*m*_ is the membrane capacitance and *λ*_v_ is the interior solution conductivity, the membrane capacitor is fully charged and the net charge of the membrane (sum of the accumulated charge on the two surfaces) is zero. As the frequency increases, *ω* ≳ *ω*_*c*_, the capacitor becomes short-circuited and draws current. This gives rise to an imbalance in the induced surface charge on the the opposite membrane surfaces; this imbalance (equivalent to an apparent membrane charge) correlates remarkably well with the viscosity frequency dependence, see Fig. 5. The only study of the effect of surface charge on the viscosity of bilayers, using mixtures of charged and neutral surfactants, has reported a significant increase [71]. The bilayers in our study are made of charge-neutral lipids, however, an apparent surface charge originates from excess mobile charges (ions) at the membrane surfaces. The mechanisms underlying the viscosity dependence on strain-rate and field frequency need further systematic exploration.

**FIG. 5.**
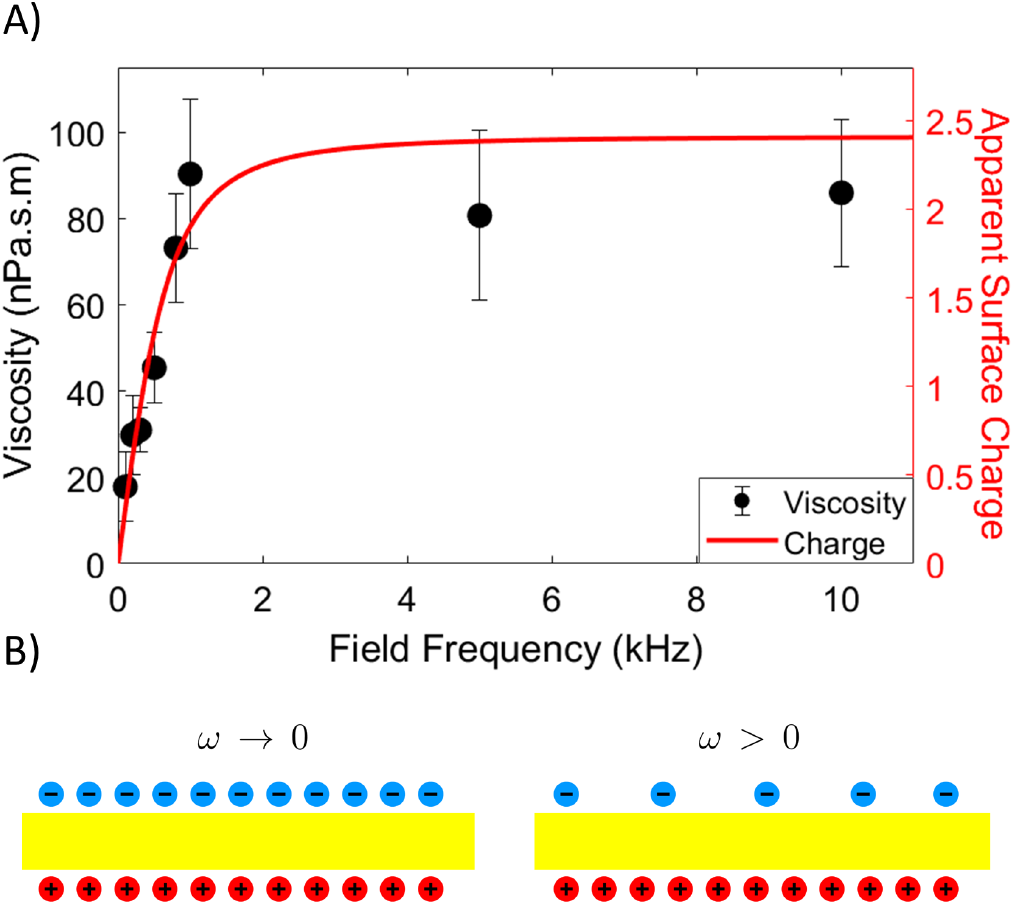
A) Membrane viscosity increases with field frequency. Data for SOPC, *E*_0_ = 8 kV/m. Same behavior is exhibited by the imbalance in the surface charges accumulated at the opposing membrane surfaces computed from Eq. 7 in SI. B) Schematic illustration of the membrane (yellow) and free charges (blue and red) under an external electric field. The membrane capacitor is fully charged at low frequencies but becomes short-circuited at higher frequencies giving rise to an imbalance of the induced charge.

**FIG. 6.**
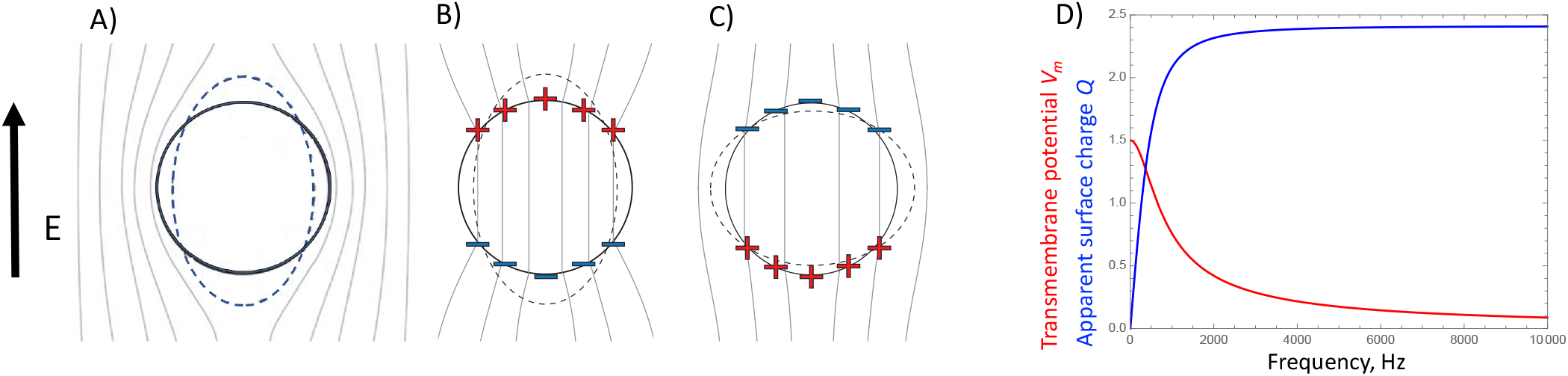
(A)-(C) Physical mechanisms of the frequency-dependent membrane polarization and vesicle dipole in an applied uniform AC field. The lines correspond to constant electric field. Upon application of an external electric field, charges accumulate on the two sides of the membrane setting up a potential difference, i.e., the membrane acts as a capacitor. (A) At low frequencies, *ω* ≪ *ω*_*c*_, the membrane capacitor is fully charged, the induced charge density on the two membrane surfaces is the same but of opposite sign. (B) and (C) At intermediate frequencies, *ω* > *ω*_*c*_, it is short-circuited and there is charge imbalance between the inner and outer membrane surfaces *Q* = *εE*_0_*Q*_0_ cos *θ*. (B) If the enclosed solution is more conducting than the suspending medium, *Λ* > 1, vesicle is pulled into an prolate ellipsoid. (C) The polarization is reversed in the opposite case *Λ* < 1 and the vesicle deforms into an oblate ellipsoid. (D) Variation with frequency of the transmembrane potential (red) and apparent charge at the pole (blue).

The ease of implementation, high-throughput, minimal experimental equipment and effort as well as robustness make this technique easy to adopt in every lab. The method can also be applied to obtain interfacial properties of lipid monolayers using deformation of droplets. We envision that our approach will become a standard tool for characterization of membrane fluidity that will help address questions of biological and engineering importance such as mechanics of excitable cells and synthetic cell design.

## MATERIALS AND METHODS

### Vesicle Preparation

Giant unilamellar vesicles (GUVs) are formed from lipids and polymer such as palmitoyloleoylphosphatidylcholine (POPC), dioleoylphosphatidylcholine (DOPC), oleoylmyristoylphosphatidylcholine (OMPC), cholesterol (Chol), stearoyloleoylphosphatdylcholine (SOPC), dipalmitoylphosphatidylcholine (DPPC) and poly(butadiene)-*b*-poly(ethylene oxide) diblock copolymers, PBd_*x*_-*b*-PEO_*y*_. The lipids and diblock copolymer were purchased from Avanti Polar Lipids (Alabaster, AL) and Polymer Source Inc. (Montreal, Canada), respectively. The multi-component vesicles made of DOPC:DPPC:Chol were fluorescently marked with 0.1 mol% of Liss Rhod PE. The lipid vesicles were produced using the electroformation method [72]. The stock solutions of 12 mM lipid in choloroform are diluted to 5 mM from which 10 *µ*l of the solution is spread on the conductive sides of the ITO slides (Delta technologies, USA). The slides are stored in vacuum for 2-4 hours to evaporate all the organic solvents. The two slides are then sandwiched with a 2 mm thick teflon spacer and the electroformation chamber is filled with 40 mM sucrose solution in 0.3 mM of NaCl. The chamber is connected to a signal generator (Agilent, USA) for 2 hours at 50 Hz and voltage 1.5 V at 60°C, which ensures that all lipids are above their main phase transition temperatures. The harvested vesicles are diluted in isotonic glucose solution without salt. 3 independent GUV batches for every lipid composition were analyzed. Polymer vesicles were produced from spontaneous swelling method. Initially, 50 *µ*l of 6-10 mg/ml (in chloroform) polymer solution was dissolved in 200-300 *µ*l of chloroform in a 20 ml vial. Polymer films were formed from evaporation by blowing with a nitrogen stream while swirling the solution inside. Afterwards, the vials were dried under vacuum for 2-4 hours. The polymer films were hydrated in the suspending solutions (40 mM sucrose solution in 0.3 mM NaCl) and placed at 60 C in an oven for 18-24 hours.

### Electrodeformation

The electrodeformation experiments are conducted in the electrofusion chamber (Eppendorf, Germany). The chamber is made from Teflon with two 92 *µ*m cylindrical parallel electrodes 500 *µ*m apart. The field is applied using a function generator (Agilent 3320A, USA). The function generator is controlled using a custom built MATLAB (Mathworks, USA) progam. This gives a precise control over the strength and duration of applied electric fields [73].

### Optical microscopy and imaging

The vesicles are visualized using a phase contrast microscope (A1 Axio Observer, Zeiss, Germany) with 63x objective 0.75 NA (air). Imaging is performed using Photron SA1.1 camera. The image acquisition rate for electrodeformation recordings is kept to a constant of 500-2000 fps for lipid vesicles and 60-200 fps for polymer vesicles and the shutter speed is fixed to 500 *µ*s. The time evolution of the vesicle is analyzed using a home-made image analysis software. The software uses a Fourier series to fit around the vesicle contour, 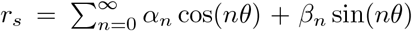. The second mode in the series is used to determine the major (*a*) and minor axis (*b*) of the deformed vesicles to evaluate 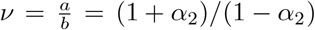. The confocal imaging was performed with Leica TCS SP8 scanning confocal microscope using a HC PL APO 40x/ NA 1.3 (oil) objective. The pinhole size during the experiment was fixed to 1 AU (Airy units). The dye was excited with a 561 nm laser (diode-pumped solid-state laser) with 1.61% (laser intensity) HyD3 detector (hybrid).

### Bending rigidity and tension measurements

Flickering spectroscopy is a popular technique to extract out membrane rigidity and tension due to its non-intrusive nature and well developed statistical analysis criteria. The details of the technique are given in [74–76]. Essentially, a time series of fluctuating vesicle contours is recorded on the focal plane. The quasi-circular contour is represented in Fourier modes, *r*(*ϕ*) = *R*(1 + ∑_*q*_ *u*_*q*_(*t*) exp(i*qϕ*). The fluctuating amplitudes *u*_*q*_ have mean square amplitude dependence on the membrane bending rigidity *κ* and the tension 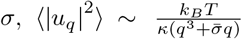, where *k*_*B*_*T* is the thermal energy (k_*B*_ is the Boltzmann constant and T is the temperature), and 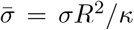. The integration time effect of the camera was reduced by acquiring images at a low shutter speed of 100-200 *µ*s. At least 5000 images were obtained for each vesicle for good statistics.

## ACKNOWLEDGMENS

P.M.V and H.A.F acknowledge financial support by NIGMS award 1R01GM140461. This research was also supported in part by the National Science Foundation under Grant NSF PHY-1748958. We thank Talbia Bint Humayun for proofreading the paper.

## Appendix A: Compendium of published values for membrane viscosity

see Table II

**TABLE II.**
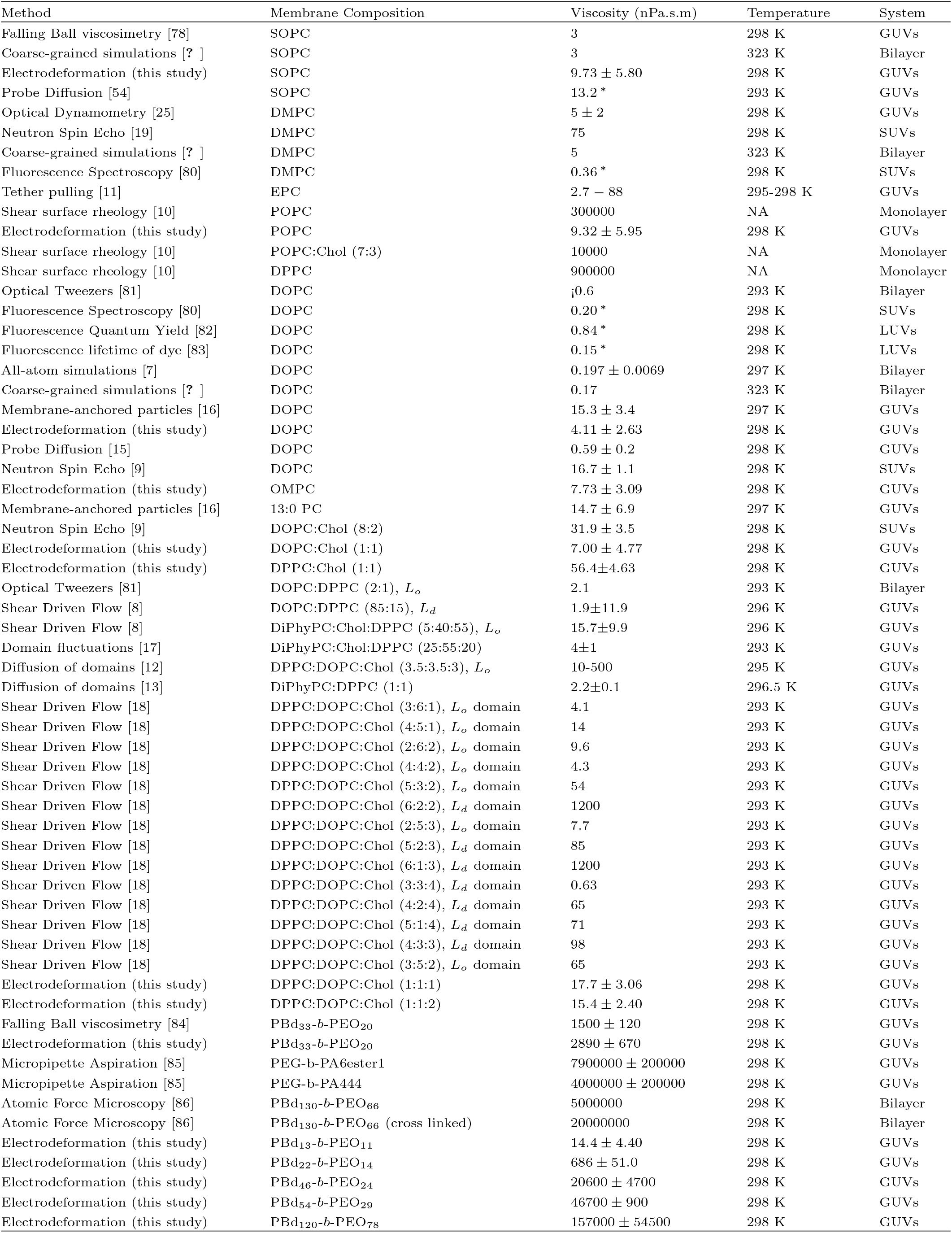
Membrane viscosity obtained using different experimental methods and simulations.* denotes membrane viscosity obtained from the bulk viscosity, *η*, using the relation *η*_*m*_ = *ηh*, where *h* is the bilayer thickness. For DMPC, SOPC and DOPC, the bilayer thickness are 3.67 nm, 4.00 nm [77] and 3.67 nm [62] respectively. *L*_*d*_ and *L*_*o*_ refer to liquid-disordered and liquid-ordered phases respectively. GUVs, LUVs and SUVs refer to Giant-Unilamellar vesicles, Large-Unilamellar vesicles and Small-Unilamellar vesicles respectively. Zero-frequency membrane viscosities obtained from the electrodeformation method developed in this study were measured at 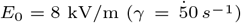.

## Appendix B: Shape evolution of quasi-spherical vesicle in a uniform electric field

### 1. Theoretical model

Let us consider a vesicle made of a charge-free lipid bilayer membrane with bending rigidity *κ*, tension *σ*_eq_, capacitance *C*_m_. The vesicle is suspended in a solution with conductivity *σ*_w_ and permittivity *ε*_w_, and filled with a different solution characterized by *σ*_v_ and *ε*_v_.

An axisymmetric stress, such as generated by uniform electric field or extensional flow, deforms the vesicle into a spheroid with symmetry axis aligned with the extensional axis. The spheroid aspect ratio is *ν* = *a/b*, where *a* is the length of the symmetry axis and *b* is the length of the axis perpendicular to the symmetry axis. For small deformations, *ν* ≲ 1.3, the shape is well approximated by

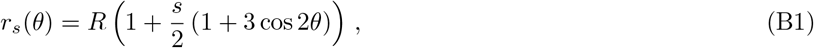

where *r*_*s*_ is the position of the surface, *R* is the initial radius of the vesicle, *s* is the deformation parameter, and *θ* is the angle with the applied field direction; *θ* = 0 and *π/*2 correspond to the pole and the equator, respectively. The ellipsoid aspect ratio is related to the deformation parameter by *ν* = (1 + *s*)*/*(1 – 2*s*).

The theory developed by *Vlahovska et al*. [45, 46, 50, 87] predicts that the deformation parameter evolution is given by the balance of imposed and membrane stresses

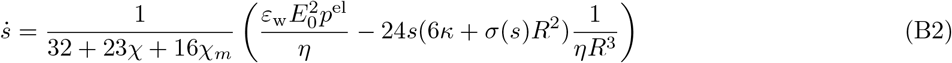

The AC field, *E*(*t*) = *E*_0_ sin(*ωt*), generates an electric stress which has two components, a steady one *p*^el^ and an oscillatory one with frequency twice the applied one

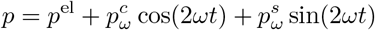

In the experiments typically 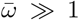 and the oscillatory component only drives very small oscillations about the deformation induced by the steady stress component.

The steady electric stress is given by

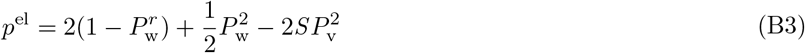

and the amplitudes of the unsteady stress are

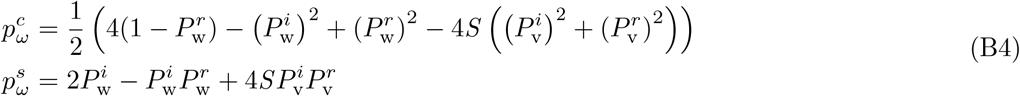

where

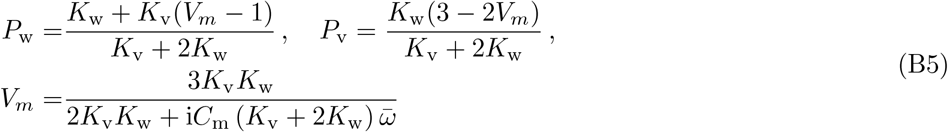

Here 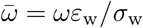 and 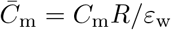 are the dimensionless frequency and membrane capacitance. 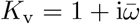 and 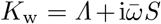 are the dimensionless complex permittivities. *S* = *ε*_v_*/ε*_w_ and *Λ* = *σ*_v_*/σ*_w_ are the ratios of permittivities and conductivities of the fluids interior and exterior to the vesicle. *P*^*r*^ and *P*^*i*^ denote the real and imaginary part of *P*, and *P* ^2^ = *PP* ^∗^, where the superscript * denotes co<u>m</u>plex conjugate. The electric stress in DC field is obtained by setting 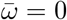 and the electric field amplitude to 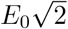.

Typically, both the inner and outer fluids are aqueous solutions with similar permittivities, *ε*_v_ ≈ *ε*_w_ = *ε*, hence *S* can be set to 1. In this case Eq. B4 reduces to

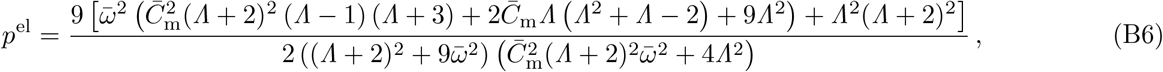

where 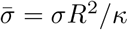. At low frequencies, 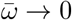, the membrane capacitor is fully charged, and *p*^el^ = 9*/*16 and we obtain Equation 1 in the main text.

The imbalance between the induced charge of the two membrane surfaces is *Q* = *εE*_0_*Q*_0_ cos *θ*, where the maximum charge is

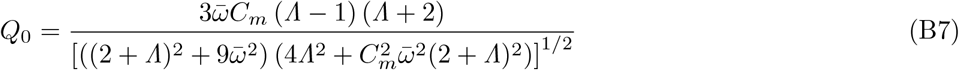

At low frequencies, 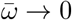 the charge imbalance vanishes.

### 2. Linear approximation of the evolution equation

Assuming constant tension, Eq. 1 in the main text can be integrated to yield

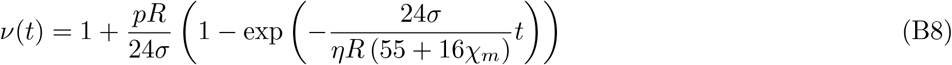

If

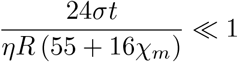

the exponent is expanded in Taylor series to yield the linear evolution, Eq. 2 in the main text.

The time limit for the linear approximation can be estimated by comparing the linear and quadratic terms in the Taylor series of the exponential term in Eq. B8. The Taylor series of the exponential function is

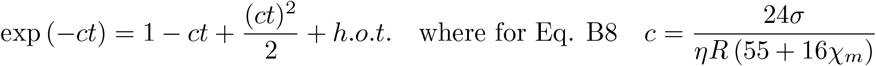

It shows that the quadratic correction becomes comparable to the linear term when

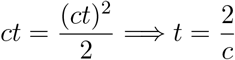

which gives the estimate for the time up to which the linear approximation is reasonable

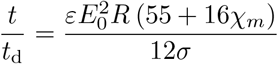

## Appendix C: Additional data

### 1. Batch reproducibility for homogeneous (DOPC) and mixed membrane compositions (DOPC:DPPC:Chol (1:1:1))

Figure 7 shows the box plot presentation for apparent membrane viscosity values obtained for the same system across three different batches of vesicles prepared from DOPC and DOPC:DPPC:Chol (1:1:1). The zero charge or frequency membrane viscosity data is given in the main text.

**FIG. 7.**
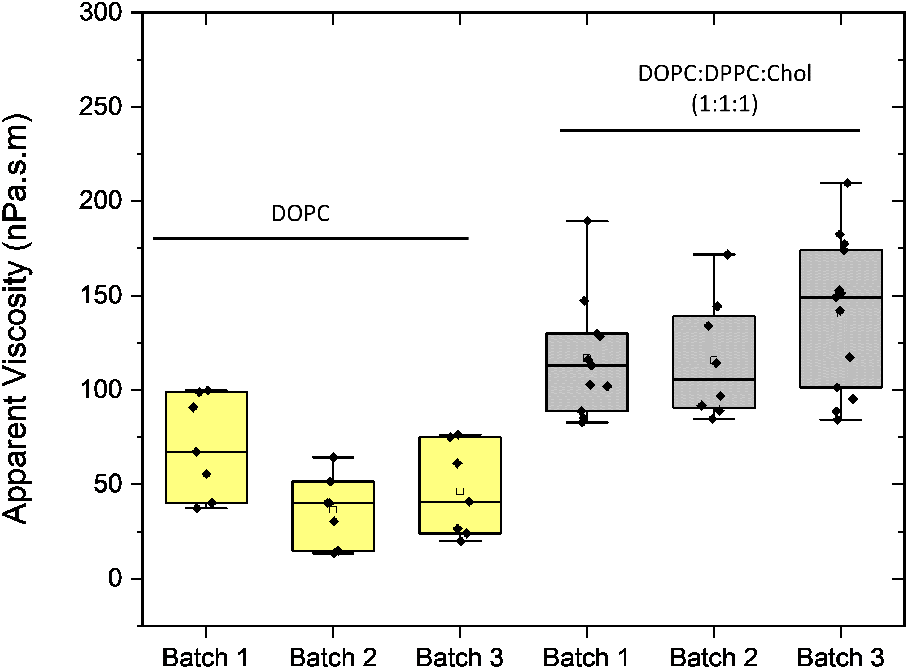
Method reproducibility for the same system across different batches: Membrane viscosity for DOPC and DOPC:DPPC:Chol (1:1:1) for three different batches of prepared vesicles. The applied frequency and field strength are 1 kHz and 10 kV/m for respectively. For DOPC:DPPC:Chol (1:1:1), the applied frequency is 2 kHz and and applied field strength is 6 kV/m. The solid symbols show measurements on individual vesicles. The box-plot represents the standardized distribution of data based on five numbers minimum value, first quartile (Q1), median, third quartile (Q3), and maximum value. The open square represents the mean value..

### 2. Deformation curves of bilayers at different field strength and frequency

Figure 8A represents deformation curves of POPC vesicles at different field strength (6-10 kV/m) at frequency 1 kHz. In Figure 8B the data is re-plotted again in rescaled time with *t*_d_. The inverse of *t*_d_ can be expressed as shear rate, 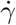 which in this case ranges from, 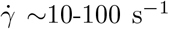. The collapse of the data on single curve indicates that the deformation rate of POPC bilayers are not affected at a given shear rate and they exhibit Newtonian rheology.

**FIG. 8.**
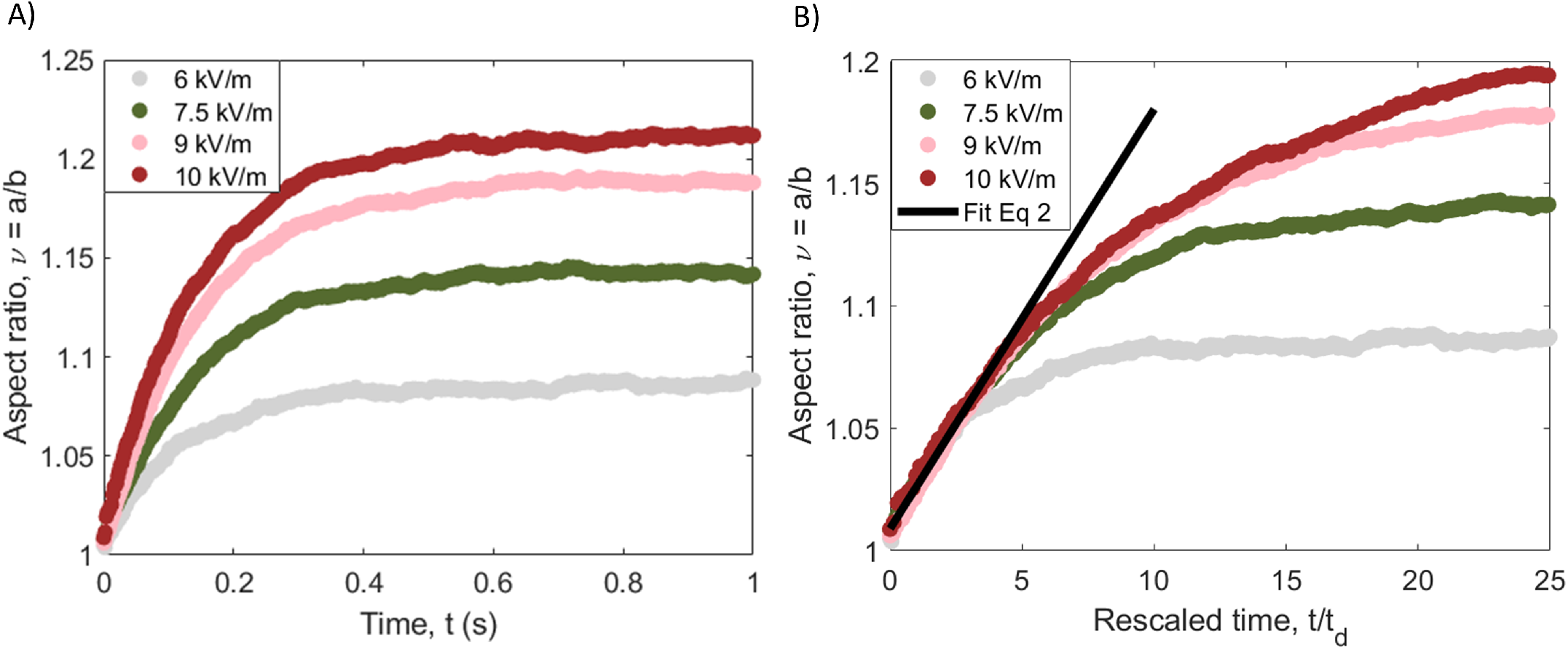
(A) Deformation curves for a POPC vesicle (R= 30.1 *µ*m) exposed to fields of different amplitudes (at 1 kHz). (B) The initial slope of the data in (A) re-plotted as a function of the re-scaled time *t/t*_d_ yields an apparent membrane viscosity *η*_*m*_ = 2.63 ± 0.41 × 10^−7^ Pa.s.m.

Eq. 2 in the main text shows that the slope depends on *t*_d_, which depends on the field amplitude *E*_0_, and *p*^el^, which depends on frequency. Hence to isolate the viscosity, one needs to plot the deformation data as a function of time rescaled as *t/t*_d_*p*^el^, see Figure 9.

**FIG. 9.**
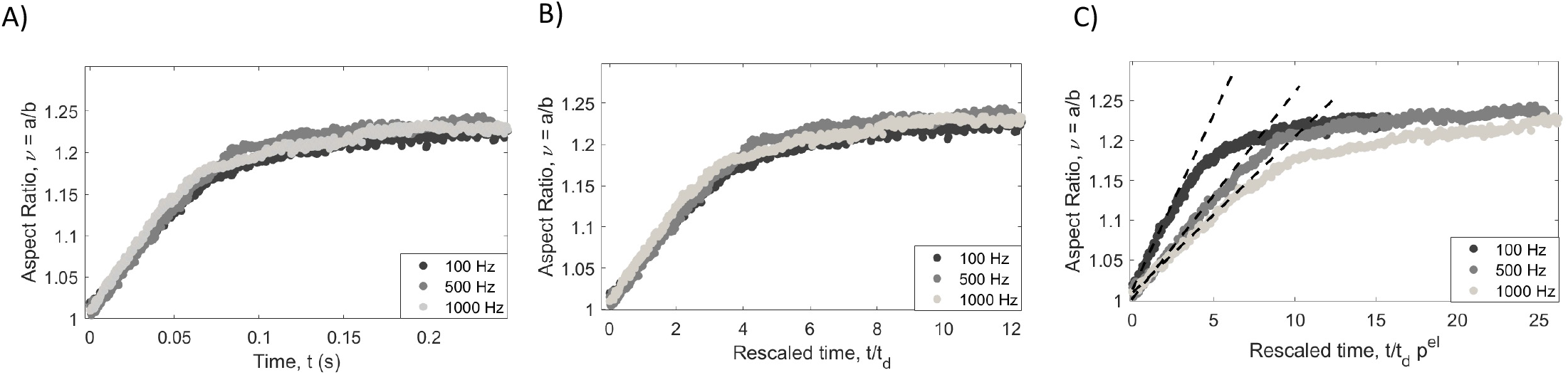
(A) Deformation curves for a POPC vesicle (R= 14.7 *µ*m) exposed to fields of different frequency but same field amplitude *E*_0_ = 8 kV/m. (B) The initial slope of the data in (A) re-plotted as a function of the re-scaled time *t/t*_d_, see Figure 9. The electric stress *p*^el^ increases with frequency but the slope in (B) remains the same indicating that apparent surface viscosity also increases. (C) Indeed, when data are plotted vs *t/t*_d_*p*^el^ the slope decreases with increasing frequency yielding the frequency dependence of the apparent viscosity. Extrapolation to zero frequency gives the membrane viscosity

### 3. Newtonian and Non-Newtonian bilayers

Figure 10 shows flow curve experiment (membrane viscosity vs imposed shear rate) for different compositions. Unsaturated phospholipids, for example DOPC in 10A and B, are known to assemble into bilayers in the liquid disordered, *L*_*d*_ phase. Our data shows that at different shear rate, 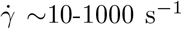, or E-field ranging from 5-40 kV/m, the membrane viscosity of *L*_*d*_ bilayers remains unchanged. Therefore, DOPC bilayers are 2D Newtonian behavior. In the next case, see 10C, we analyze, fully saturated lipids, DPPC, mixture with Cholesterol at 1:1 molar ratio. DPPC has a phase transition temperature temperature at 314 K. Adding Cholesterol to the bilayers results in the disruption of the gel or solid phase of the bilayer to *L*_*o*_ state which has a larger extent of orientational order than *L*_*d*_. Our results show that *L*_*o*_ are also Newtonian in nature. Next we move to mixed systems exhibiting phase separation. 1:1 DOPC:DPPC demonstrated phase separation with intricate network of finger-like domains indicative of gel or solid phase. Interestingly, the flow curve experiments,10D and E, show Non-Newtonian characteristics with shear thinning behavior (undergoing flow with a decreasing viscosity with increasing shear rate). Polymer vesicles assembled from di-block copoylmers such as PBd_*x*_-*b*-PEO_*y*_ also show a shear thinning behaviour with increasing shear rate. 10F and G gives two examples of PBd_*x*_-*b*-PEO_*y*_ bilayers where the higher *M*_*w*_ polymersomes, PBd_33_-*b*-PEO_20_, show a much stronger shear thinning behaviour. The viscosity of PBd_33_-*b*-PEO_20_ decreases by a 100% from 622 ± 70 × 10^−8^ Pa.s.m to 311 ± 62 × 10^−8^ Pa.s.m between shear rate 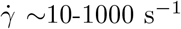. PBd_13_-*b*-PEO_11_ viscosity decreases from 8.72 ± 2.22 × 10^−8^ Pa.s.m to 5.6 ± 0.5 × 10^−8^ Pa.s.m between similar shear rates.

**FIG. 10.**
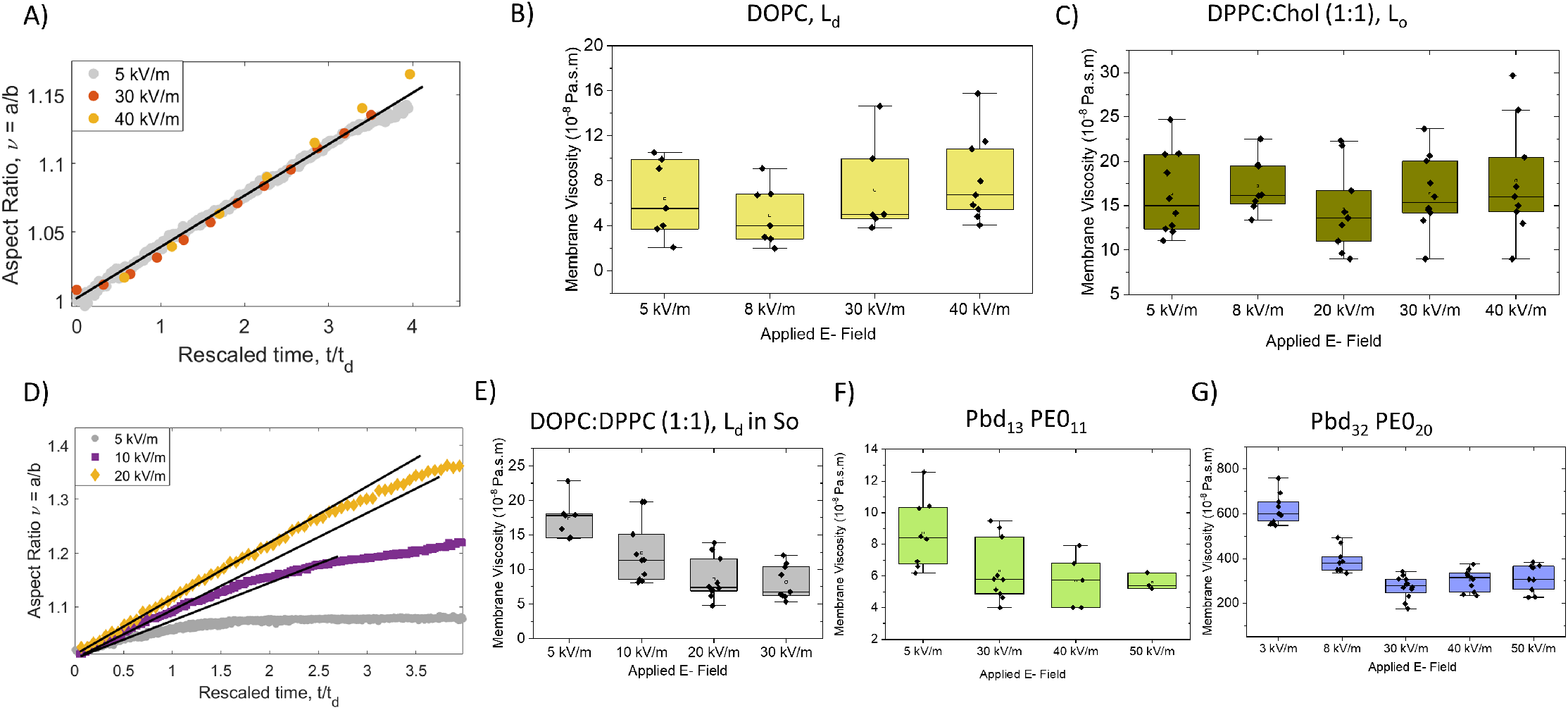
Shear viscosity of different bilayer compositions at different electric field (3-50 kV/m) or shear rates 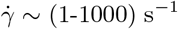 at a fixed frequency 1 kHz. A) Deformation case of DOPC bilayer at different field strengths. The slope remains same regardless of the applied shear rate indicating 2D Newtonian fluids. B) Viscosity obtained at different field strengths for Newtonian cases B) DOPC C) DPPC:Chol (1:1) bilayers. D) Deformation case of DOPC:DPPC (1:1) bilayer at different field strength. The slope increases as the field strength or shear rate is increased. This behavior is represented by shear thinning fluids. B) Viscosity obtained at different field strengths for non-Newtonian cases E) DOPC:DPPC (1:1) F) PBd_13_-*b*-PEO_11_ G) PBd_32_-*b*-PEO_20_.

### 4. Bending rigidity values from Flickering Spectroscopy and capacitance measurements for electrodeformation method

The method for flickering spectroscopy is detailed in [75, 76]. Here, we summarize the electrodeformation method to extract out membrane capacitance. The procedure follows the original approach developed by *Salipante et al*. [88]. The vesicle shape morphology with conductivity ratio *Λ >* 1 is always prolate. However, for *Λ <* 1, the conductivity of the outer solution is higher than the vesicle solution and the aspect ratio/deformation parameter *s* (*ω*) is positive at low frequencies that is prolate shape. As the frequency increases, the vesicles becomes less prolate and adopts a spherical shape at a certain frequency. Above this critical frequency, the vesicles adopt an oblate shape. The critical frequency can be approximated as:

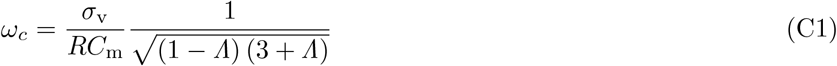

Hence, the membrane capacitance can be determined from the experimentally measured critical frequency based on prolate-oblate transition with a frequency sweep [88]. The measured bending rigidity and capacitance values are summarized in Table III.

**TABLE III.**
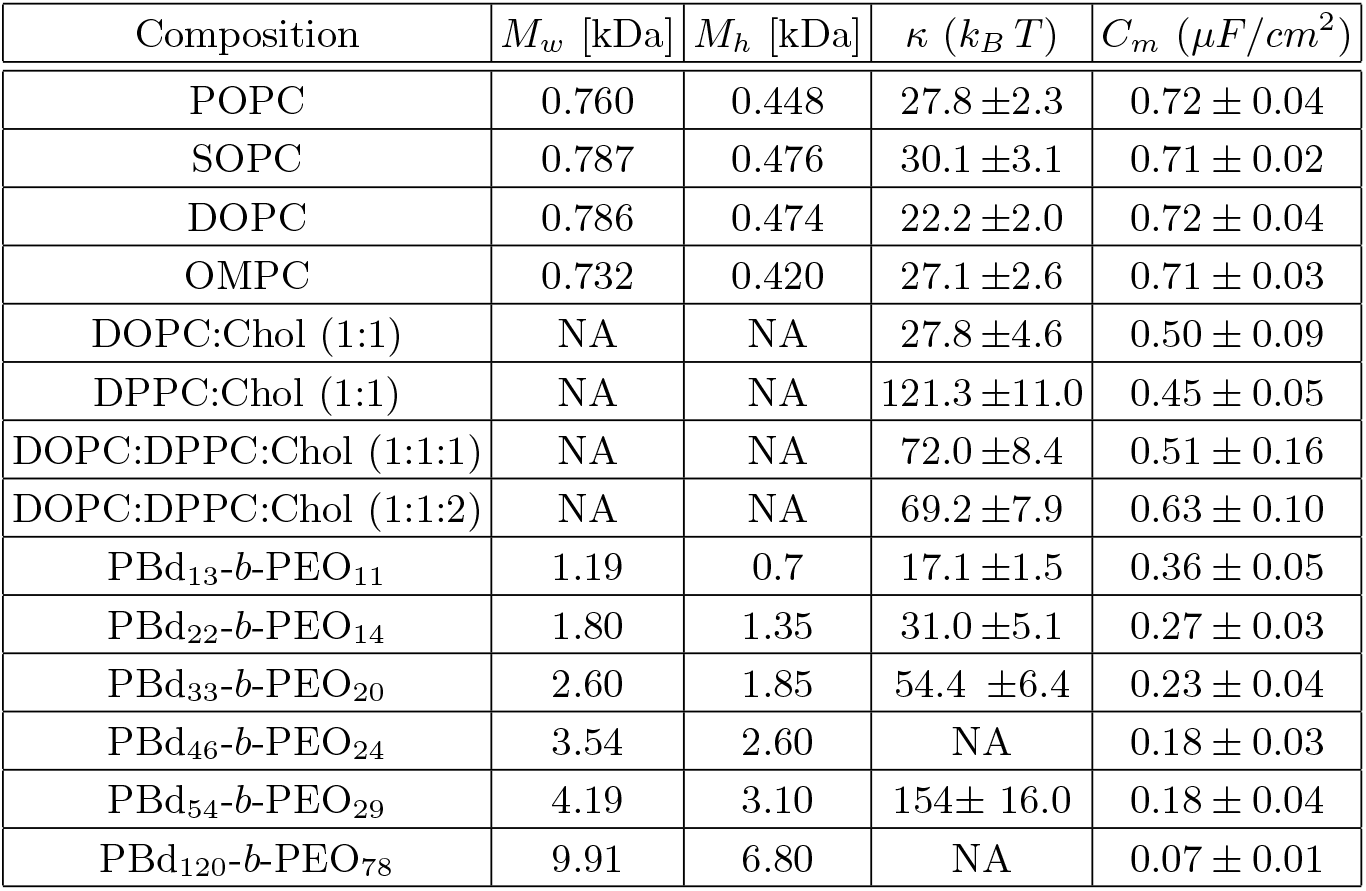
Membrane bending rigidity and capacitance of phospholipids, polymers PBd_*x*_-*b*-PEO_*y*_ and mixed system of DOPC:DPPC:Chol at 25 *°*C determined in this study. Bending rigidity was measured with flickering spectroscopy and membrane capacitance was measured with the electrodeformation method. *M*_*w*_ and *M*_*h*_ refer to the total and hydrophobic molecular weight, respectively. NA means not available.

**TABLE IV.** Membrane viscosities and values of a dye diffusion coefficient (DiC18) for the DOPC:DPPC:Chol ternary system. The values in brackets indicate lipid molar ratios (first column) and the number of measured vesicles (third column). All the experiments were performed at 25.0 *°*C. *L*_*d*_ and *L*_*o*_ denote liquid disordered and liquid ordered, respectively. The diffusion coefficient were taken from [55]

| Multi-component | Phase state | $\eta_m$ [nPa.s.m] | D [$\mu\text{m}^2/\text{s}$ [55]] |
| --- | --- | --- | --- |
| DOPC | $L_d$ | $4.11 \pm 2.63$ (20) | $6.30 \pm 0.13$ |
| DPPC:Chol (1:1) | $L_o$ | $56.4 \pm 4.63$ (25) | $0.48 \pm 0.06$ |
| DOPC:DPPC:Chol (1:1:2) | $L_o$ | $15.4 \pm 2.40$ (25) | $1.85 \pm 0.13$ |
| DOPC:DPPC:Chol (1:1:1) | $L_d$ | $17.7 \pm 3.06$ (18) | $2.50 \pm 0.20$ |
| DOPC:Chol (1:1) | $L_d$ | $7.00 \pm 4.77$ (25) | $3.25 \pm 0.25$ |

## Appendix D: Rheology of block copolymer PBd_33_-*b*-PEO_20_

The copolymers PBd_33_-*b*-PEO_20_ was obtained from Polymer Source Inc. (Montreal, Canada). Figure 11A and B demonstrate the molecular formula and morphology of the melt at room temperature. The melt appears to be like a pasty wax. The shear rheology response was characterized using Anton MCR rheometer 302 for rheological testing. A parallel plate (PP 25 mm) fixture was used for the measurements for low viscosity measurements. The parallel plate gap width was kept 0.7 to *mm*. 0.5 gm of the sample was loaded on the rheometer. The bulk shear viscosity, 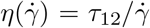, was determined from the measured shear stress, *τ*_12_, from imposed shear rates in the range from 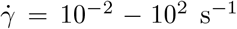. As shown in Figure 11C, the strain was chosen to be 1 % to ensure that the sample was measured in linear viscoelastic regime. Figure 11D represents the viscosity at different shear rates. The block co polymer demonstrates consistent shear thinning, and total lack of zero-shear regime. The power law obtained in -0.51. This power law behavior is consistent with other published data in literature [89].

**FIG. 11.**
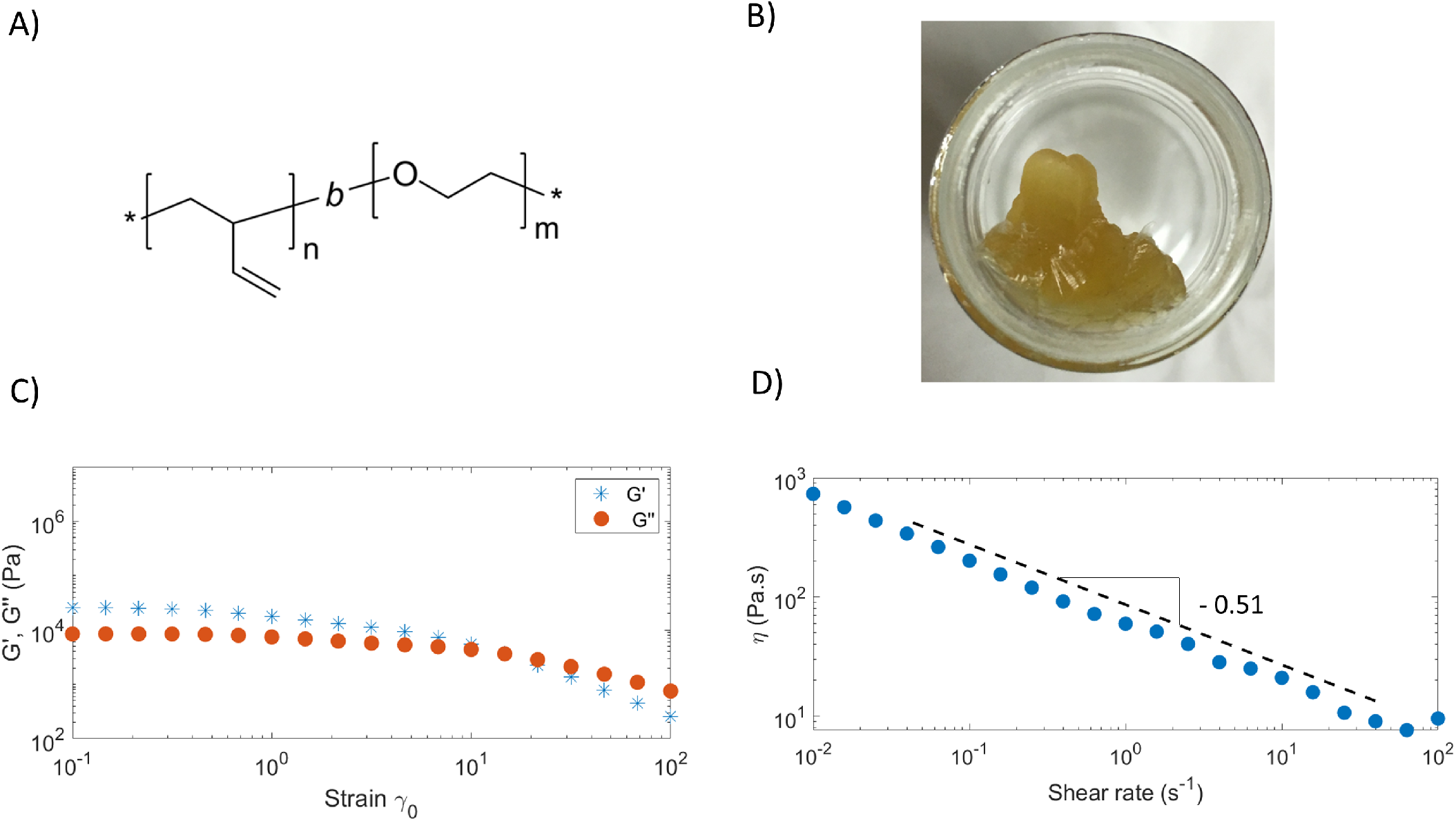
A) Molecular structure of PBd_*n*_-*b*-PEO_*m*_ diblock co-polymer. B) The appearance of the polymer melt at room temperature. C) Determination of loss and storage modulus at different strain to determine the linear viscoelastic strain limit. D) The flow curve experiment with bulk viscosity vs shear rate.

